# The transcriptional repressor HIC1 regulates intestinal immune homeostasis

**DOI:** 10.1101/083873

**Authors:** Kyle Burrows, Frann Antignano, Michael Bramhall, Alistair Chenery, Sebastian Scheer, Vladimir Korinek, T. Michael Underhill, Colby Zaph

**Affiliations:** The Biomedical Research Centre, University of British Columbia, Vancouver, British Columbia, V6T 1Z3, Canada; Department of Pathology and Laboratory Medicine, University of British Columbia, Vancouver, British Columbia, V6T 2B5, Canada; The Department of Cellular & Physiological Sciences, University of British Columbia, Vancouver, British Columbia, V6T 1Z3, Canada; Department of Cell and Developmental Biology, Institute of Molecular Genetics, Academy of Sciences of the Czech Republic, 142 20 Prague 4, Czech Republic; Infection and Immunity Program, Monash Biomedicine Discovery Institute, School of Biomedical Sciences, Monash University, Clayton, VIC, 3195, Australia; Department of Biochemistry and Molecular Biology, School of Biomedical Sciences, Monash University, Clayton, VIC, 3195, Australia

## Abstract

The intestine is a unique immune environment that must respond to infectious organisms but remain tolerant to commensal microbes and food antigens. However, the molecular mechanisms that regulate immune cell function in the intestine remain unclear. Here we identify the POK/ZBTB family transcription factor Hypermethylated in cancer 1 (HIC1, ZBTB29) as a central component of immunity and inflammation in the intestine. HIC1 is specifically expressed in immune cells in the intestinal lamina propria (LP) in the steady state and mice with a T cell-specific deletion of HIC1 have reduced numbers of T cells in the LP. HIC1 expression is regulated by the Vitamin A metabolite retinoic acid, as mice raised on a Vitamin A-deficient diet lack HIC1-positive cells in the intestine. HIC1-deficient T cells overproduce IL-17A in vitro and in vivo, and fail to induce intestinal inflammation, identifying a critical role for HIC1 in the regulation of T cell function in the intestinal microenvironment under both homeostatic and inflammatory conditions.

## INTRODUCTION

The intestinal immune system is constantly interacting with non-pathogenic commensal organisms and innocuous food antigens that must be tolerated immunologically, while maintaining the ability to rapidly respond to infectious organisms and toxins. To maintain this balance, unique immune cell populations are found in the intestinal microenvironment. For example, intestinal-resident dendritic cells (DCs) that express the surface marker CD103 have been shown affect intestinal homeostasis by producing the vitamin A metabolite retinoic acid (RA) (Scott, et al. 2011). Intestinal macrophages have also been shown to play a central role in tolerance in the intestine (Smith, et al. 2010; Bain and Mowat, 2014). Specific subsets of CD4^+^ T helper (T_H_) cells, primarily FOXP3-expressing regulatory T cells (T_reg_ cells) and IL-17A-producing T_H_ cells (T_H_17 cells) are enriched in the intestinal lamina propria (LP) (Littman and Rudensky, 2010), while CD8^+^ T cells and γδ T cells are found in the intraepithelial compartment (Cheroutre, et al. 2011), acting in conjunction with IgA-producing B cells that are required to maintain tolerance to the commensal flora (Suzuki, et al., 2007). However, the molecular mechanisms that regulate intestinal immune cell function in health and disease are not fully defined.

The POZ and Kruppel/Zinc Finger and BTB (POK/ZBTB) proteins are a family of transcription factors that play critical roles in a variety of biological processes such as gastrulation, limb formation, cell cycle progression, and gamete formation (Kelly and Daniel, 2006). In the immune system, POK/ZBTB proteins are key regulators in cellular differentiation and function (Lee and Maeda, 2012). For example, B-Cell CLL/Lymphoma 6 (BCL6, ZBTB27) is critical for the development of germinal centres (GCs) following immunization or infection. BCL6 is expressed at high levels in GC B cells (Allman et al., 1996) and T follicular helper (T_FH_) cells (Nurieva et al., 2009) that together control the germinal centre response. Promyelocytic leukaemia zinc finger (PLZF, ZBTB16) is required for NKT cell development (Savage et al., 2008) and expression of PLZF has recently been shown to identify a multipotent progenitor of innate lymphoid cells (ILCs) (Constantinides et al., 2014). T helper-inducing POZ/Kruppel-Like factor (ThPOK, ZBTB7B) is a master regulator of T_H_ cell development in the thymus (Wang et al., 2008; Muroi et al., 2008). Hypermethylated in cancer 1 (HIC1, ZBTB29) is a member of this family and has been shown to regulate proliferation and p53-dependent survival in a wide range of tumours. HIC1 is epigenetically silenced through DNA methylation in various human cancers (Chen et al., 2003; Wales et al., 1995) and it has been proposed that HIC1-depdent repression of SIRT1 was critical for the function of p53 (Chen et al., 2005). However, the role of HIC1 in immune cells has not been examined.

In this study, we identify HIC1 as a regulator of intestinal immune responses under homeostatic and inflammatory conditions. We demonstrate that HIC1 is expressed in immune cells specifically in the intestinal lamina propria but not in other lymphoid or non-lymphoid tissues in the steady state. In the absence of HIC1 in T cells, we observe a significant reduction in the frequency of T cells in the LP and intraepithelial compartment, coincident with increased T_H_17 cell responses. HIC1 expression in the LP is regulated by the vitamin A metabolite retinoic acid and under inflammatory conditions in the intestine, loss of HIC1 specifically in T cells renders mice resistant to the development of intestinal inflammation, suggesting that HIC1 is required for the pathogenicity of T cells in vivo. Thus, HIC1 plays a central role in intestinal immune homeostasis and inflammation.

## RESULTS

### HIC1 expression in immune cells is restricted to the intestine

To begin to address the role of HIC1 in the immune system, we examined the expression of HIC1 in CD45^+^ leukocytes in various tissues of mice with a fluorescent reporter gene inserted in the *Hic1* locus (*Hic1*^Citrine^ mice) (Pospichalova et al., 2011). Citrine expression in CD45^+^ cells was restricted to the intestine, with the only detectable Citrine-positive cells in the lamina propria and intraepithelial space (Figure 1a). Further characterization of the leukocytes in the LP revealed that a majority of T cell receptor β (TCRβ) chain-expressing CD4^+^ and CD8^+^ T cells and TCRγδ^+^ CD8^+^ T cells in the LP expressed HIC1 (Figure 1b) and that TCRβ^+^ CD8^+^ and TCRγδ^+^ CD8^+^ intraepithelial lymphocytes (IELs) also expressed HIC1 (Figure 1c). We also found that most MHCII^+^ CD11c^+^ CD64^-^-dendritic cells (DCs) and MHCII^+^ CD64^+^ F4/80^+^ macrophages expressed HIC1 (Figure 1–figure supplement 1). However, HIC1 expression was not generalized for all lymphocytes in the LP, as B220^+^ B cells did not express HIC1 (Figure 1b). Thus, HIC1 expression in immune cells specifically identifies intestinal resident populations.

**Figure 1.**
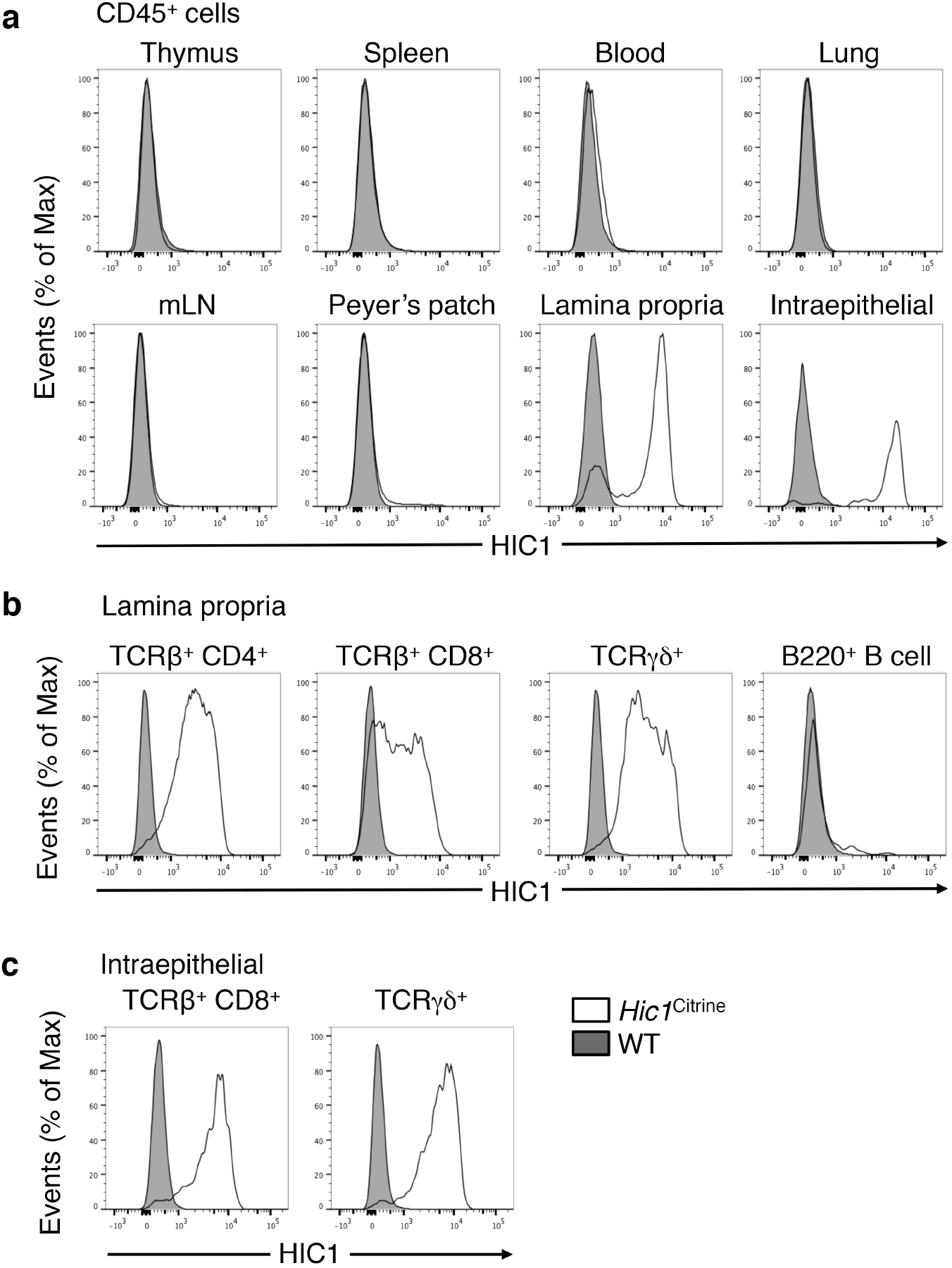
HIC1 expression in immune cells is restricted to the intestine. (a) Steady state thymus, spleen, blood, lung, mesenteric lymph node (mLN), peyer’s patch (PP) and intestinal lamina propria and intraepithelial leukocytes were analyzed by flow cytometry for *Hic1^Citrine^* reporter expression in total CD45^+^ leukocytes. (b) TCRβ^+^CD4^+^ T cells, TCRβ^+^CD8^+^ T cells, TCRγδ^+^ T cells or B220^+^ B cells were analyzed by flow cytometry for *Hic1^Citrine^* reporter expression from the intestinal lamina propria or (c) intraepithelial compartment. (a-c) Data are representative of 3 ndependent experiments.

### T cell-intrinsic expression of HIC1 regulates intestinal immune homeostasis

As HIC1 was expressed in multiple immune cell populations in the LP, we next sought to determine the cell-intrinsic functions of HIC1 by focusing on its role specifically in T cells. To do this, we crossed mice with *loxP* sites flanking the *Hic1* gene (*Hic1^fl/fl^* mice) with mice that express the Cre recombinase under control of the *Cd4* enhancer and promoter (*Cd4*-Cre mice) to generate mice with a T cell-intrinsic deletion of *Hic1* (*Hic1^ΔT^* mice). *Hic1^ΔT^* mice displayed normal thymic development (Figure 2–figure supplement 1a,b) and had normal frequencies of CD4^+^ and CD8^+^ T cells in the spleen (Figure 2–figure supplement 1c) or mesenteric lymph nodes (mLN) (Figure 2–figure supplement 1d). HIC1 also had no effect on the activation state of splenic CD4^+^ T cells, as we observed equivalent expression of CD62L and CD44 between *Hic1^f/f^* and *Hic1^ΔT^* mice (Figure 2–figure supplement 1e). Finally, HIC1 was not required for the development of FOXP3^+^ CD25^+^ CD4^+^ T cells in the spleen or mLNs (Figure 2–figure supplement 1f). Thus, HIC1 expression is dispensable for peripheral T cell homeostasis. However, we found that the frequency and number of CD4^+^ and CD8^+^ TCRβ^+^ cells in the LP (Figure 2a,b) and CD8^+^ T cells in the intraepithelial compartment (Figure 2a,c) was significantly reduced in the absence of HIC1, demonstrating that T cell-intrinsic expression of HIC1 is required for intestinal T cell homeostasis.

**Figure 2.**
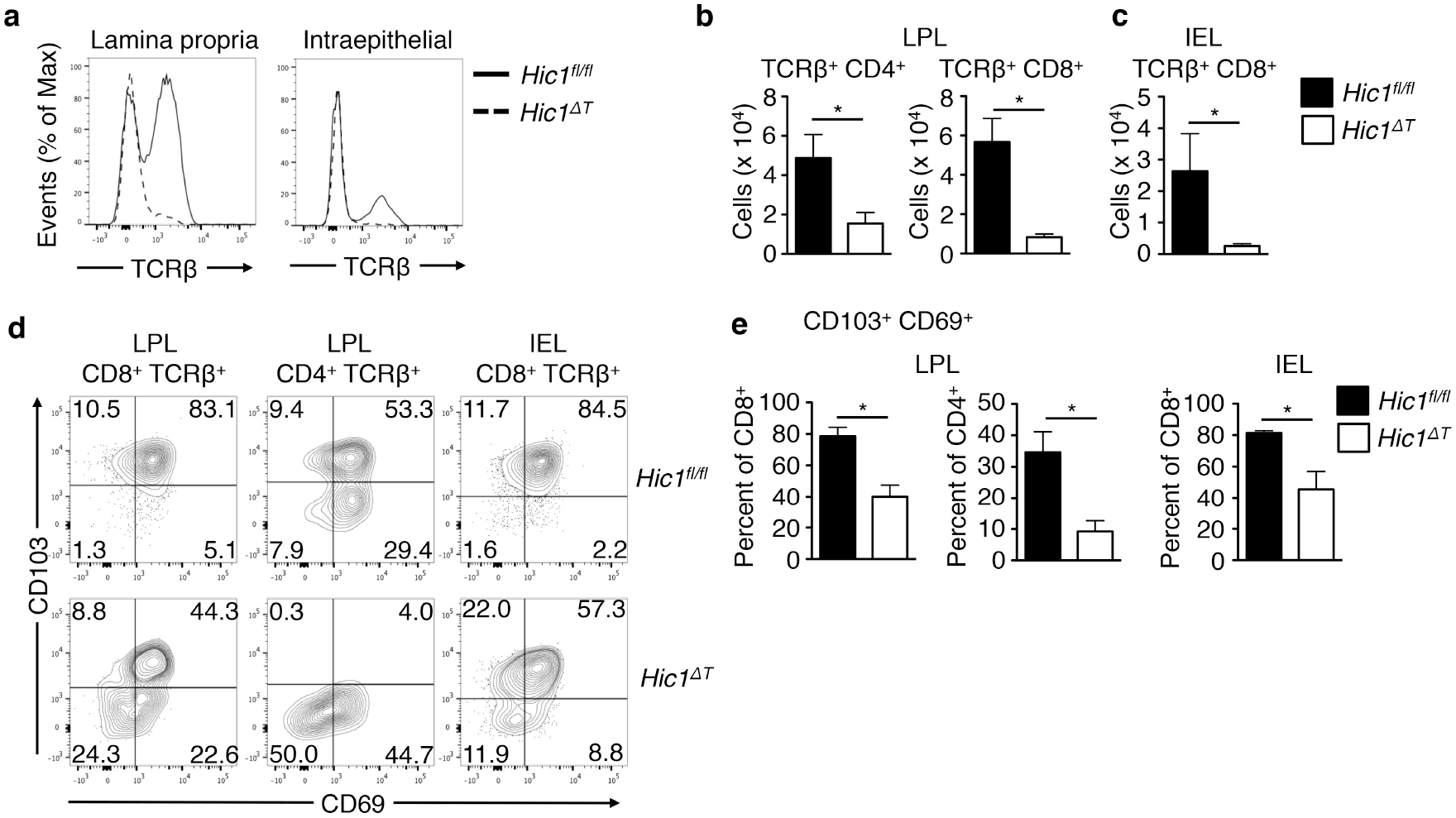
T cell-intrinsic expression of HIC1 regulates intestinal T cell homeostasis. Intestinal lamina propria (LPL) and intraepithelial (IEL) lymphocytes were isolated from *Hic1^fl/fl^* or *Hic1^ΔT^* mice. (a) Frequency of TCRβ^+^ T cells and (b,c) total number of TCRβ^+^CD4^+^ and TCRβ^+^CD8^+^ T cells were analyzed by flow cytometry. (d) Flow cytometry analysis and (e) quantification of CD103 and CD69 surface marker expression from IEL and LPL TCRβ^+^CD4^+^ and TCRβ^+^CD8^+^ T cells. (a,d) Data are representative of 3 independent experiments. (b,c,e) Data are pooled from 3 independent experiments (n=6-7 per group). * *P* < 0.05. Error bars indicate SEM.

Lamina propria lymphocytes (LPLs) and intraepithelial lymphocytes (IELs) display an activated, memory phenotype that includes the markers CD69 and CD103, cell surface molecules that are important for retention in the intestinal microenvironment. CD69 has been shown to negatively regulate expression of sphingosine-1-phosphate receptor (S1PR1), which must be downregulated to establish tissue-residency (Matloubian et al., 2004; Skon et al., 2013), while CD103 binds to the epithelial cell-expressed E-cadherin and is required for maintenance of intestinal T cells (Schön et al., 1999). We found that HIC1-deficient CD4^+^ and CD8^+^ T cells in the LP and intraepithelial compartments expressed significantly reduced levels of CD69 and CD103 (Figure 2d,e). Strikingly, there was an almost complete loss of CD103^+^ CD69^+^ CD4^+^ T cells in the LP (Figure 2d,e). Thus, the reduced frequency and number of CD4^+^ and CD8^+^ LPLs and IELs in *Hic1^ΔT^* mice is associated with reduced expression of CD69 and CD103.

### HIC1 is a negative regulator of IL-17A production by CD4^+^ T cells

In the intestinal LP, CD4^+^ T_H_17 and T_reg_ cells are found at higher frequencies than other T_H_ cell subsets. Analysis of the CD4^+^ LPLs showed that although frequency of FOXP3^+^ T_reg_ cells were equivalent between *Hic1^fl/fl^* and *Hic1^ΔT^* mice, the frequency of RORγt^+^ IL-17A-expressing T_H_17 cells in *Hic1^ΔT^* mice was significantly increased (Figure 3a,b). These results suggest that in addition to regulating T cell numbers in the intestine, HIC1 deficiency in T cells had a specific effect on T_H_17 cell differentiation. To directly examine the role of HIC1 in T_H_ cell differentiation, we stimulated T_H_ cells that were purified from the spleen and peripheral lymph nodes (pLNs) of naïve *Hic1^fl/fl^* and *Hic1^ΔT^* mice (Figure 3c) under diverse polarizing conditions. HIC1 deficiency had no effect on the development of T_H_1, T_H_2 or TGF-β-induced T_reg_ (iT_reg_) cells (Figure 3–figure supplement 1). Consistent with our *in vivo* results, we observed a significant increase in the production of IL-17A by HIC1-deficient T_H_ cells activated under T_H_17 cell-promoting conditions, with heightened frequency and mean fluorescence intensity of IL-17A^+^ cells (Figure 3d), resulting in a significant increase in the levels of secreted IL-17A (Figure 3e), which was also associated with increased *Il17a* expression in HIC1-deficient T_H_17 cells (Figure 3f). Retroviral transduction of HIC1 into differentiating T_H_17 cells resulted in a complete inhibition of IL-17A production (Figure 3g,h). Taken together, these results suggest that HIC1 is a cell-intrinsic negative regulator of IL-17A production during T_H_17 cell differentiation.

**Figure 3.**
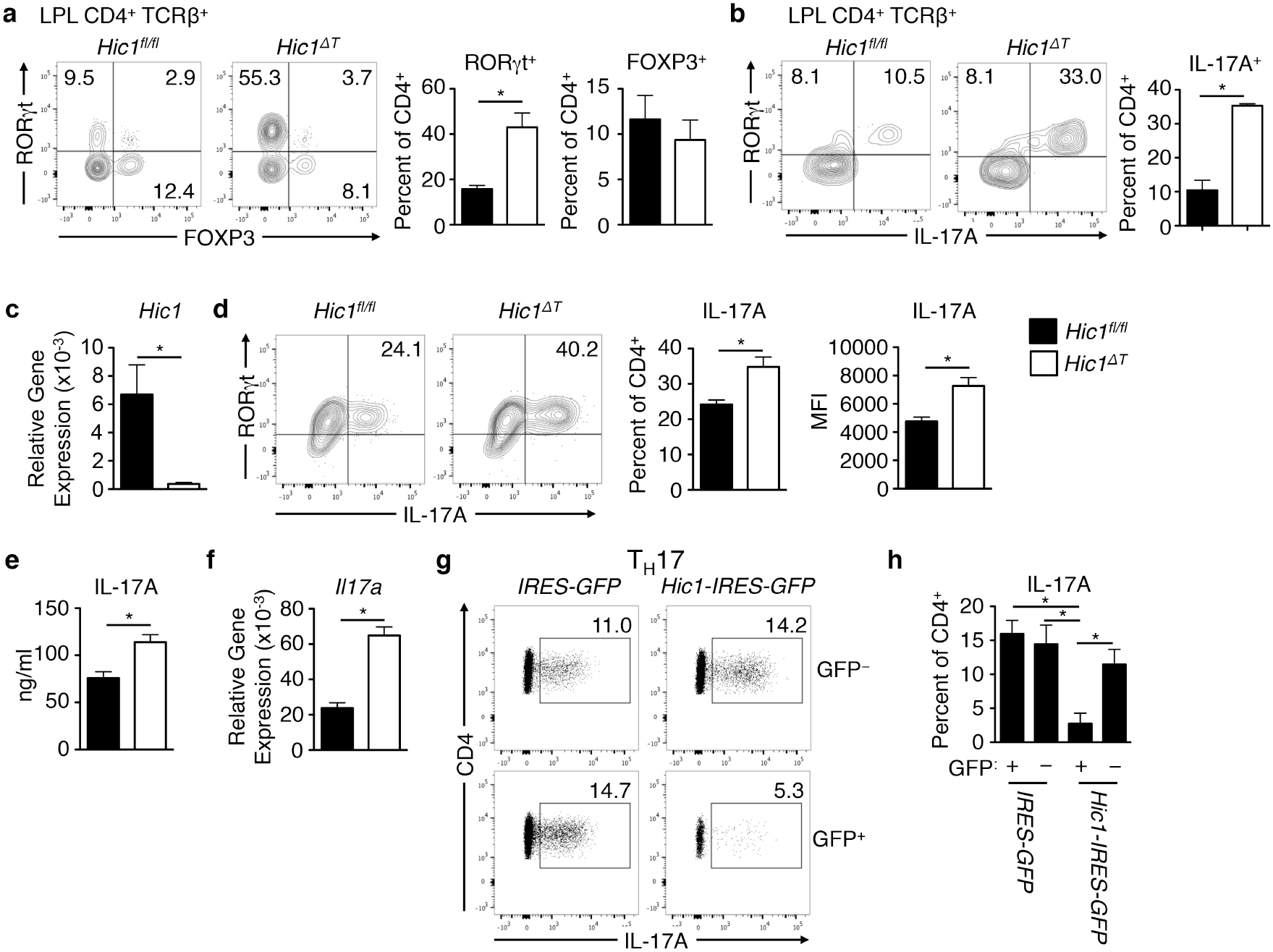
HIC1 is a negative regulator of IL-17A production by CD4^+^ T cells. (a,b) Steady-state intestinal lamina propria (LPL) CD4^+^ T cells were isolated from *Hic1^fl/fl^* or *Hic1^ΔT^* mice and analyzed by flow cytometry for intracellular expression of RORγt, FOXP3 and IL-17A. Data are pooled from 3 independent experiments (n=5-6 per group). *Hic1^fl/fl^* or *Hic1^ΔT^* splenic CD4^+^ T cells were activated under T_H_17 cell-polarizing conditions and analyzed for: (c) *Hic1* mRNA expression by quantitative RT-PCR, (d) intracellular RORγt and IL-17A frequency and mean fluorescence intensity (MFI) by flow cytometry, (e) IL-17A protein production by ELISA, and (f) *Il17a* mRNA expression by qRT-PCR. Data are pooled from 4 independent experiments (n=4-8 per group). (g-h) Analysis of IL-17A expression from T_H_17 cell-polarized cells that were retrovirally transduced with MigR1-IRES GFP (empty vector) or MigR1-*Hic1*-IRES GFP expression vectors. GFP^+^ indicated successful retroviral infection. (h) Data are pooled from 3 independent experiments (n=3 per group). * *P* < 0.05. Error bars indicate SEM.

### Retinoic acid regulates expression of HIC1 in T cells in vitro and in vivo

The vitamin A metabolite RA plays a critical role in intestinal immune homeostasis (Brown and Noelle, 2015a). Previous studies have shown that mice raised on a diet lacking vitamin A (Vitamin A-Deficient (VAD) diet) have defects in T_H_ cell activation and intestinal migration, resulting in an overall impairment in T cell-driven immune responses in the intestine (Iwata et al., 2004; Johansson-Lindbom et al., 2005; Hall et al., 2011). To test whether RA influenced HIC1 expression in intestinal T_H_ cells, we raised *Hic1*^Citrine^ mice on a VAD diet. We found that LP T_H_ cells from VAD mice failed to express HIC1 (Figure 4a), suggesting that HIC1 expression in T_H_ cells in the LP is dependent upon the presence of RA. Consistent with this, addition of RA to T_H_ cells isolated from spleen or lymph nodes of mice activated *in vitro* with antibodies against CD3 and CD28 resulted in an upregulation of HIC1 at the mRNA and protein levels (Figure 4b,c). Analysis of HIC1 expression using *Hic1*^Citrine^ mice further demonstrated that treatment of activated T_H_ with RA led to increased HIC1 expression (Figure 4d). Expression of HIC1 was dependent upon T_H_ cell activation, as addition of RA to T_H_ cells in the absence of T cell receptor stimulation had no effect on the expression of *Hic1* mRNA (Figure 4e). Thus, these results identify HIC1 as an RA-responsive gene in activated T_H_ cells.

**Figure 4.**
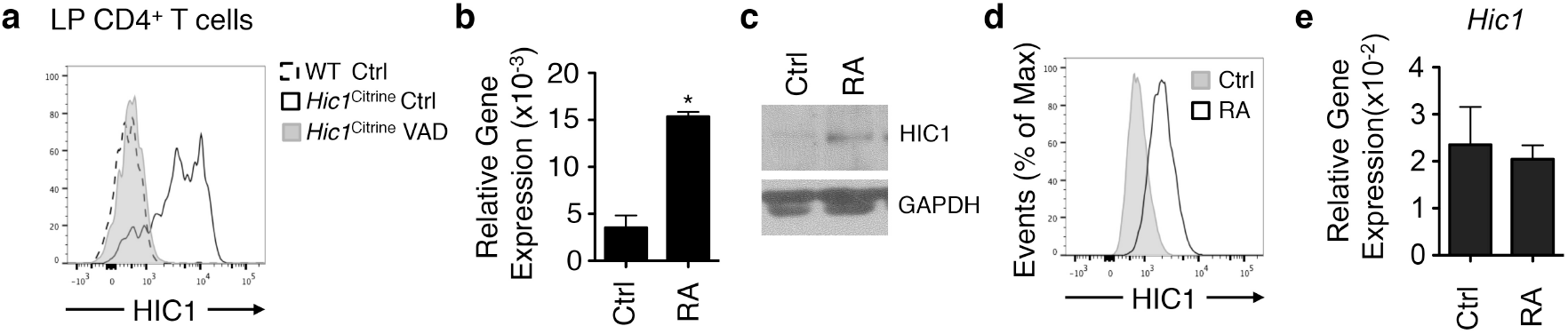
Retinoic acid regulates expression of HIC1 in CD4^+^ T cells. (a) HIC1 reporter expression in intestinal lamina propria (LP) CD4^+^ T cells from *Hic1^Citrine^* mice fed a control diet, *Hic1*^Citrine^ mice fed a vitamin A deficient (VAD) diet, and controls fed a control diet was analyzed by flow cytometry. Data are representative of 3 independent experiments. (b-d) Splenic T_H_ cells were activated with antibodies against CD3 and CD28 in the presence or absence of 10 nM RA and HIC1 levels were measured by (b) quantitative RT-PCR (c) western blot and (d) HIC1 reporter expression. (b) Data are pooled from 2 independent experiments (n=4 per group). (c and d) Data are representative of 2 independent experiments. (e) Expression of *Hic1* in naïve splenic CD4^+^ T cells that were treated with 10 nM RA for 16 hours was analyzed by qRT-PCR. Data are pooled from 2 independent experiments (n=4 per group). * *P* < 0.05. Error bars indicate SEM.

### HIC1 is dispensable for expression of intestinal homing receptors

RA is critical for migration of immune cells to the intestine (Iwata et al., 2004) through the upregulation of intestinal homing molecules including α4β7 integrin and CCR9 (Johansson-Lindbom et al., 2005). Our results showing that *Hic1^ΔT^* mice had reduced numbers of intestinal T cells could be due to reduced expression of intestinal homing molecules, resulting in impaired intestinal migration. However, we failed to observe any reduction in expression of α4β7 and CCR9 on T_H_ cells in the presence or absence of HIC1 following activation in the presence of RA (Figure 5a). Further, we found that T_H_ cells isolated from the Peyer’s patches or mLNs of *Hic1^ΔT^* mice were not deficient in expression of CCR9 (Figure 5–figure supplement 1). These results demonstrate that other HIC1-dependent mechanisms such as reduced expression of CD69 and CD103 were responsible for the paucity of T_H_ cells in the LP and that HIC1 is dispensable for RA-dependent induction of intestinal homing molecules.

**Figure 5.**
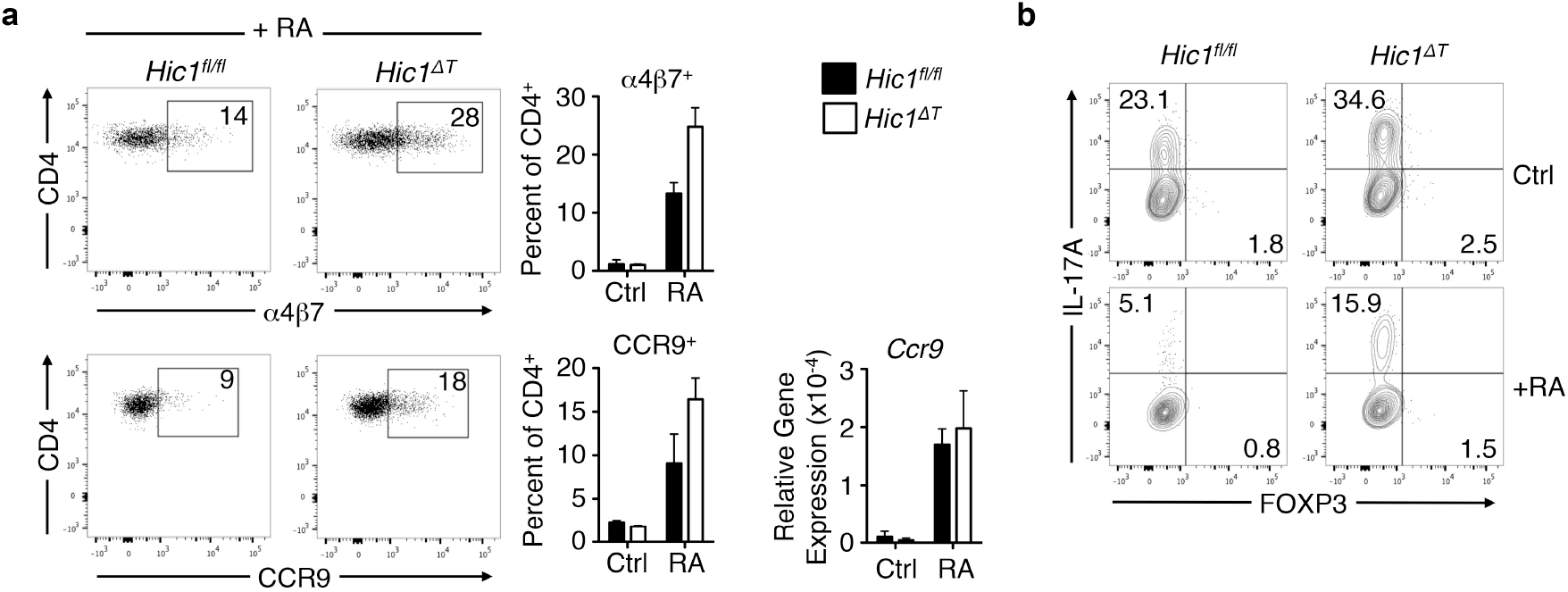
HIC1 is not required for all RA-mediated effects on CD4^+^ T cells. (a) Splenic T_H_ cells were activated with antibodies against CD3 and CD28 in the presence of 10 nM RA; α4β7^+^ and CCR9 levels were measured by flow cytometry and quantitative RT-PCR. Data are pooled from 2 independent experiments (n=4 per group). (b) Splenic CD4^+^ T cells were activated under T_H_17 cell-polarizing conditions in the presence or absence of 10 nM RA and analyzed for intracellular IL-17A and FOXP3 by flow cytometry. Data are representative of 3 independent experiments. * *P* < 0.05. Error bars indicate SEM.

### HIC1 is not required for inhibitory effects of RA on T_H_17 cell differentiation

Several studies have demonstrated that RA can negatively affect the differentiation of T_H_17 cells (Mucida et al., 2007; Hill et al., 2008; Schambach et al., 2007) although the precise mechanisms remain unclear. Based on our data showing heightened IL-17A production by HIC1-deficeint T_H_17 cells in vitro and in vivo, we hypothesized that expression of HIC1 would be required for the RA-dependent reduction in T_H_17 cell differentiation. In contrast to our expectations, addition of RA to either HIC1-sufficient or -deficient T_H_ cells led to a significant reduction in the expression of IL-17A (Figure 5b). Thus, although HIC1 is upregulated by RA in T_H_ cells and *Hic1*-deficient T_H_17 cells express increased levels of IL-17A in vitro and in vivo, HIC1 dispensable for RA-dependent regulation of T_H_17 cell responses in vitro.

### HIC1 regulates T cell-mediated inflammation in the intestine

We next examined whether dysregulated T_H_ cell responses observed in naive *Hic1^ΔT^* mice had any effect on the development of intestinal inflammation. Despite the reduced numbers of T_H_ cells in the LP of *Hic1^ΔT^* mice and the dysregulated production of IL-17A by *Hic1*-deficient T_H_17 cells, we failed to observe any significant differences in intestinal architecture between naïve *Hic1^fl/fl^* or *Hic1^ΔT^* mice (Figure 6a, left panels). After induction of intestinal inflammation with intraperitoneal injection of a monoclonal antibody against CD3 (Rutz et al., 2015; Esplugues et al., 2011; Merger et al., 2002), *Hic1^ΔT^* mice displayed less intestinal inflammation compared to control *Hic1^fl/fl^* mice (Figure 6a,b). Although we observed fewer CD4^+^ T cells in the intestine of treated *Hic1^ΔT^* mice (Figure 6c), we did observe an increase in the number of CD4^+^ T cells in the LP of treated *Hic1^ΔT^* mice compared to naïve *Hic1^ΔT^* mice (Figure 2a,b), further demonstrating that intestinal migration is not completely impaired in the absence of HIC1. Analysis of cytokine production by T_H_ cells from the LP of anti-CD3 treated mice identified an increased frequency of IL-17A-producing T_H_ cells without any changes in the frequency of IFN-γ-producing cells (Figure 6c), similar to our results under steady-state conditions. Thus, T cell-intrinsic expression of HIC1 modulates inflammation in the intestine, potentially by negatively regulating IL-17A production.

**Figure 6.**
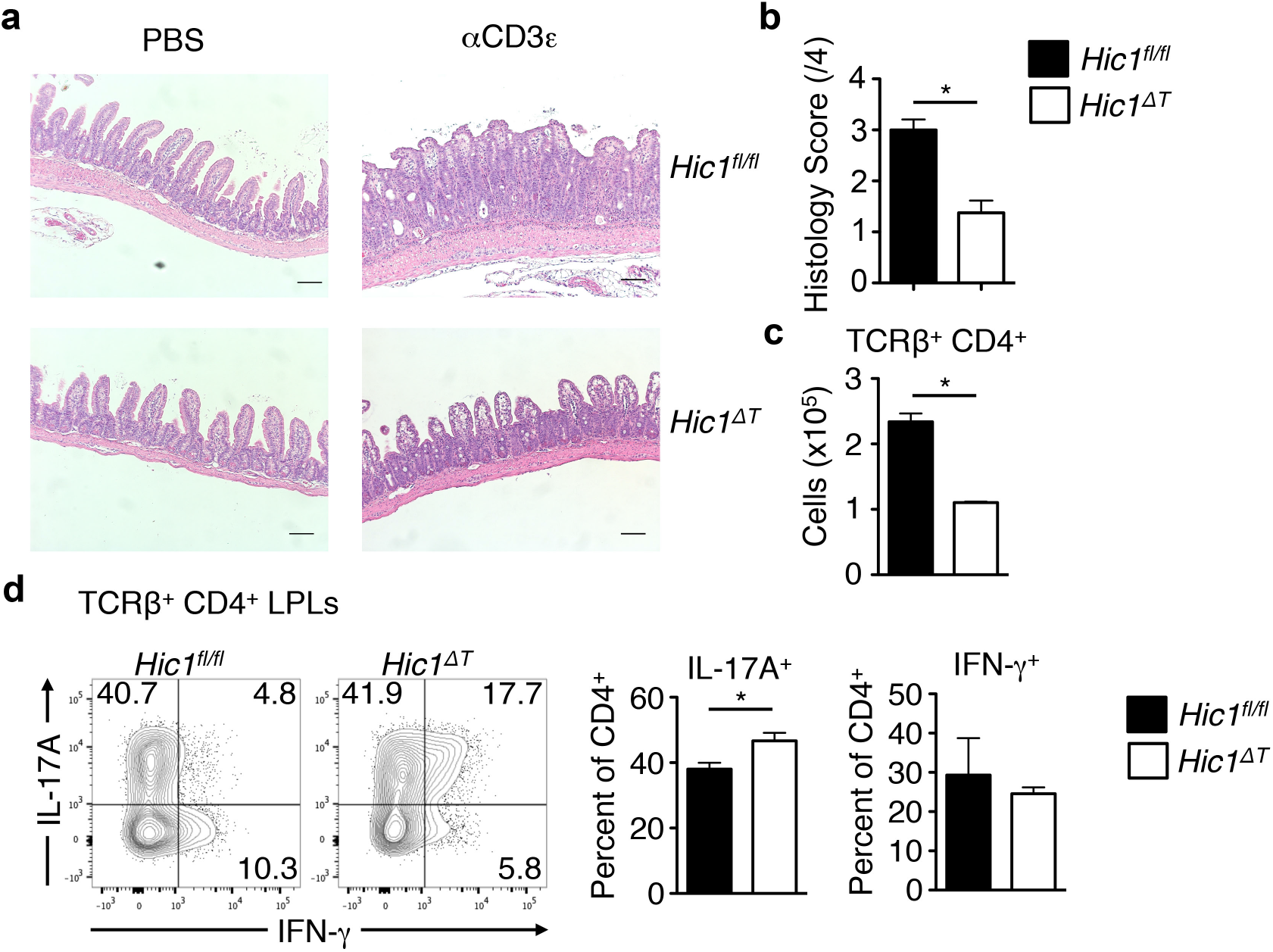
HIC1 is required for the development of intestinal inflammation. Mice received anti-CD3ε antibody by intraperitoneal injection. (a) At 48 hours post-injection, mice were sacrificed and analyzed for intestinal tissue pathology and inflammatory infiltrate by H&E staining. Scale bar represents 100μm. (b) Histological scores. (c) Total numbers of intestinal lamina propria (LPL) TCRβ^+^CD4^+^ T cells were quantified by flow cytometry. (d) Intracellular expression of IL-17A and IFN-γ from LPL CD4^+^ T cells were analyzed by flow cytometry. (a-d) Data are from 3 independent experiments (n=8-9 per group). * *P* < 0.05. Error bars indicate SEM.

To directly assess the cell-intrinsic role of HIC1 in intestinal inflammation, we employed a T cell transfer model of intestinal inflammation (Powrie et al., 1993). We adoptively transferred naïve CD4^+^ CD25^−^ CD45RB^high^ T cells isolated from either *Hic1^fl/fl^* or *Hic1^ΔT^* mice into *Rag1*^−/−^ mice. *Rag1*^−/−^ mice that received control T cells from *Hic1^fl/fl^* mice began losing weight around 4 weeks post-transfer (Figure 7a) and showed significant intestinal inflammation by 6 weeks post-transfer (Figure 7b). Strikingly, we found that *Rag1*^−/−^ mice that received T cells from *Hic1^ΔT^* mice continued to gain weight and did not develop severe intestinal inflammation (Figure 7a-c). Associated with the reduced disease, there were significantly fewer CD4^+^ T cells in the intestinal LP (Figure 7d). Furthermore, consistent with our results under homeostatic conditions as well as following α-CD3 treatment, we found that the absence of HIC1 in T cells resulted in heightened production of IL-17A with no significant effect on the frequency of IFN-γ-positive cells (Figure 7e). Thus, HIC1 expression in T cells is critically required for the development of intestinal inflammation, possibly by limiting expression of IL-17A.

**Figure 7.**
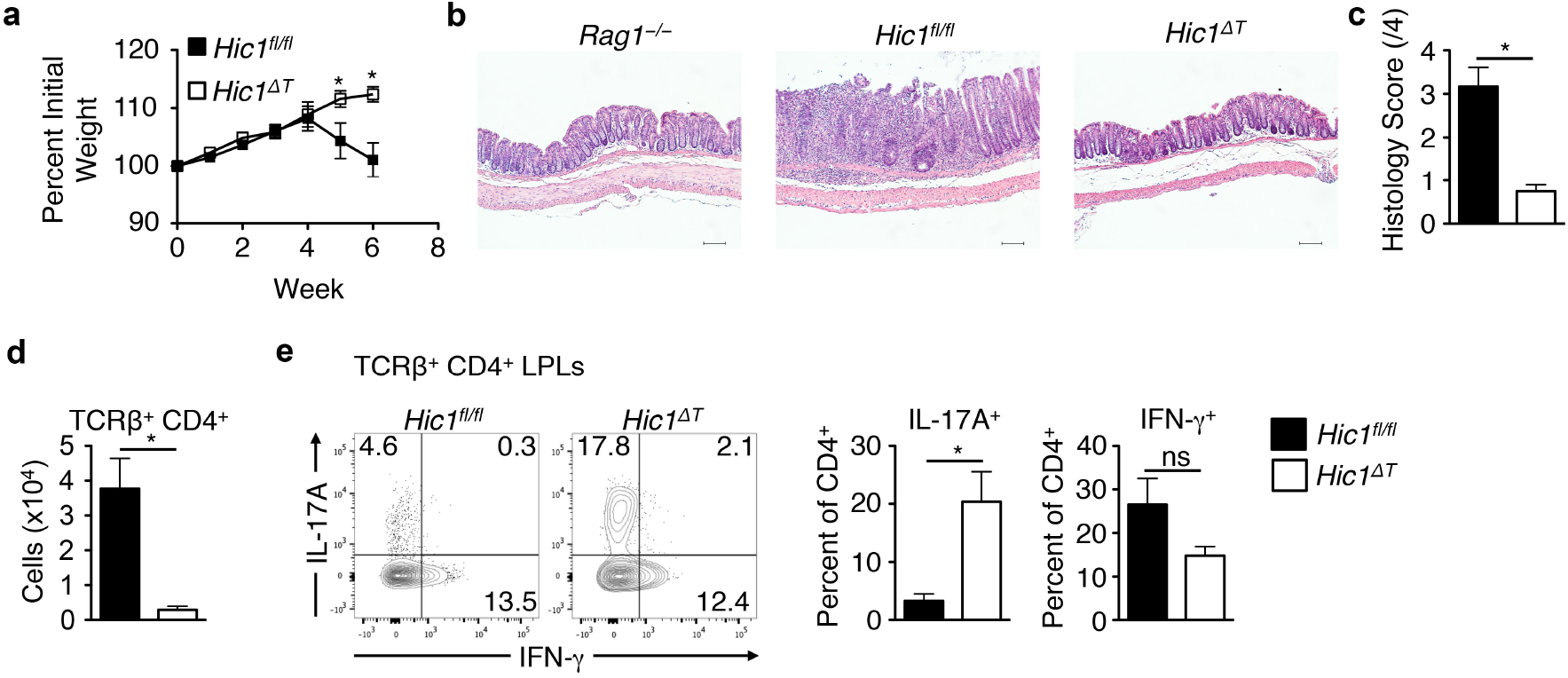
HIC1-deficient T cells fail to promote intestinal inflammation following adoptive transfer into *Rag1^−/−^* mice. CD4^+^CD25–CD45RB^hi^ naive T cells (4 × 10^5^) from *Hic1^fl/fl^* or *Hic1 δ^T^* mice were transferred into *Rag1^−/−^* mice and monitored for colitis. (a) Weight loss (percentage of initial weight) was calculated for each mouse over 6 weeks. (b) At 6 weeks post-transfer, mice were sacrificed and analyzed for intestinal tissue pathology and inflammatory infiltrate by H&E staining. Scale bar represents 100μm. (c) Histological scores. (d) Total number of TCRβ^+^CD4^+^ T cells isolated from intestinal lamina propria (LPL). (e) Intracellular expression of IL-17A and IFN-γ from LPL CD4^+^ T cells were analyzed by flow cytometry. (a-e) Data are pooled 2 of 4 independent experiments (*n* = 4–8 per experiment). Statistics compare *Rag1^−/−^* mice that received *Hic1^fl/fl^* T cells to those that received *Hic1^ΔT^* T cells. * *P* < 0.05. Error bars indicate SEM.

### HIC1 is required to limit STAT3 signaling in T_H_17 cells

In addition to directly repressing target genes, HIC1 has been shown to negatively regulate gene expression by several mechanisms including the recruitment of co-repressors such as the CtBP, NuRD and polycomb repressive complex 2 (PRC2) complexes to target genes to mediate gene repression (Van Rechem et al., 2010; Boulay et al., 2012). Further, HIC1 has also been shown to indirectly repress transcription by binding to transcriptional activators and preventing binding to target genes. HIC1 has been shown to interact with and inhibit DNA binding of transcription factors such as T cell factor-4 (TCF4), β-Catenin and STAT3 (Valenta et al., 2006; Lin et al., 2013; Hu et al., 2016). As HIC1-deficient T_H_17 cells produce heightened levels of the STAT3 target gene IL-17A (Durant et al., 2010), we hypothesized that increased STAT3 activity was associated with HIC1 deficiency. We first examined the levels of IL-6-induced active phosphorylated STAT3 (pSTAT3) in HIC1-sufficient and -deficient T_H_17 cells by flow cytometry. We found that loss of HIC1 had no effect on the frequency of pSTAT3-positive T_H_ cells (Figure 8a), suggesting that HIC1 did not affect upstream STAT3 activation. However, co-immunoprecipitation studies in T_H_17 cells of either native HIC1 (Figure 8b) or retrovirally-transduced FLAG-tagged HIC1 (Figure 8c) demonstrated that HIC1 and STAT3 interacted in T_H_17 cells. Thus, our results suggest that HIC1 limits T_H_17 cell differentiation by binding to STAT3, inhibiting its DNA binding and transcriptional activation. Consistent with this hypothesis, we found increased STAT3 binding to the *Il17a* promoter in the absence of HIC1 (Figure 8d), whereas there was no difference in binding at an irrelevant site (*Il5* promoter). Taken together, our results identify an important role for HIC1 in limiting expression of IL-17A in T_H_17 cells by inhibition of STAT3 DNA binding.

**Figure 8.**
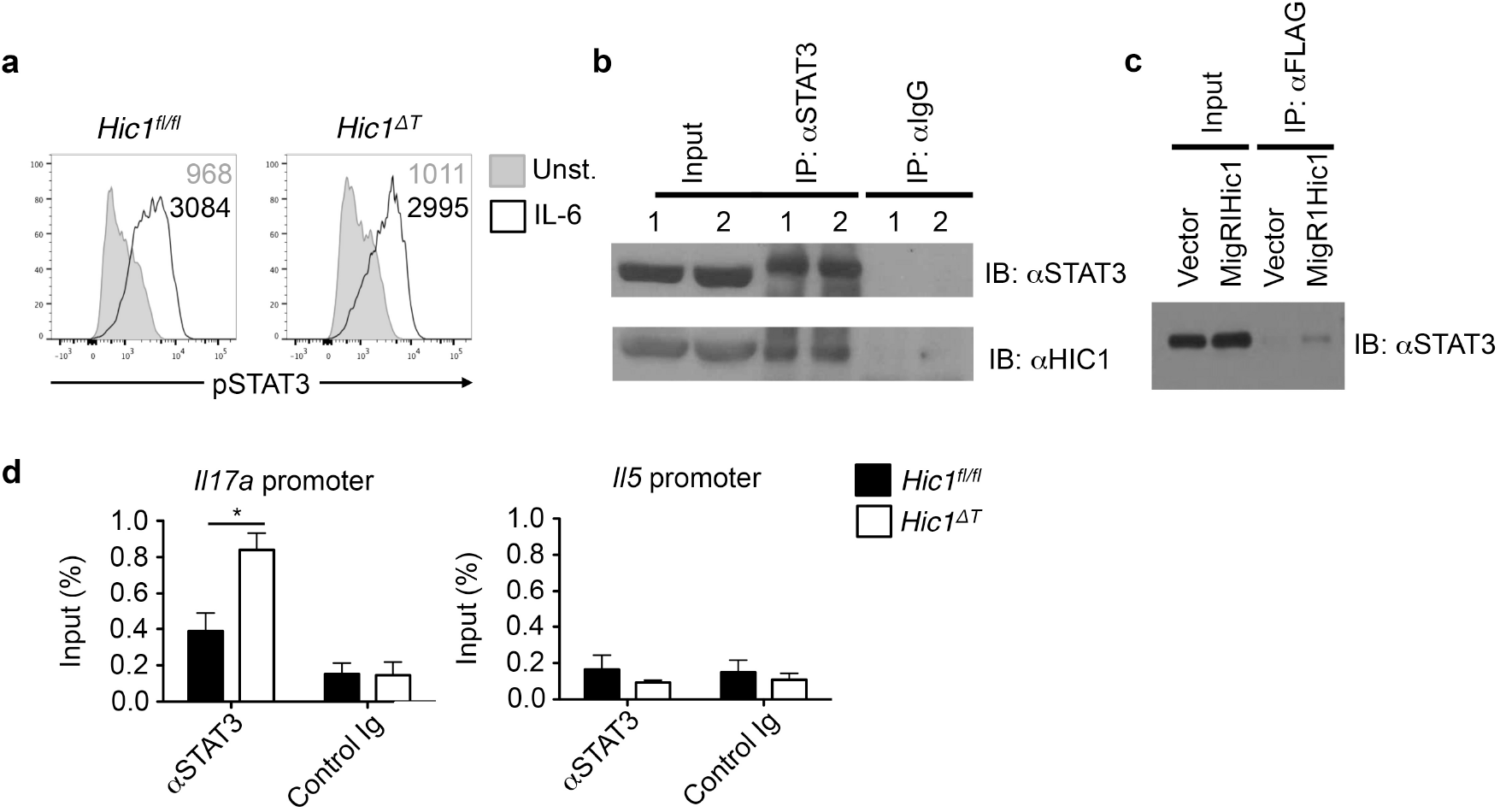
HIC1 regulates STAT3 signaling in T_H_17 cells. CD4^+^ T cells were activated under T_H_17 cell-polarizing conditions. (a) Flow cytometric analysis of STAT3 phosphorylation from T_H_17 cells restimulated with or without IL-6 for 15 minutes. Numbers represent mean fluorescence intensity. Data are representative of 2 independent experiments. (b) Inputs, anti-STAT3 immunoprecipitates (IP), and normal rat serum IP from T_H_17 cells were immunoblotted with anti-STAT3 and anti-HIC1 antibodies. 1 and 2 represent duplicate T_H_17 cultures. Data are representative of 2 independent experiments. (c) Inputs and anti-FLAG IP from T_H_17 cells retrovirally transduced with MigR1 (empty) or MigR1 FLAG-HIC1 vectors were immunoblotted with an anti-STAT3 antibody. Data are representative of 2 independent experiments. (d) Chromatin Immunoprecipitation (ChIP) analysis of STAT3 binding to the *Il17a* and *Il5* promoters in T_H_17 cells. Data are from 3 independent experiments (n=6 per group). * *P* < 0.05. Error bars indicate SEM.

### Discussion

We have identified the transcriptional factor HIC1 as a novel regulator of intestinal immune homeostasis. In the steady state, HIC1 is specifically expressed in immune cells of the intestinal LP and intraepithelial niches in an RA-dependent manner, and mice with a T cell-intrinsic deletion of HIC1 have a dramatic decrease in the number of T cells in intestinal tissues. HIC1 deficiency leads to increased expression of IL-17A, possibly through the loss of STAT3 inhibition. Under inflammatory conditions, HIC1 expression in T cells is required for intestinal pathology, identifying HIC1 as a potential therapeutic target to treat intestinal inflammation.

Our results suggest that the micronutrient RA regulates HIC1 expression in immune cells in the intestine. In support of this, two unrelated genome-wide expression screens for genes in T_H_ cells regulated by RA signaling have identified HIC1 as an RA/RARα-dependent gene, although no functional analyses were carried out (Kang et al., 2011; Brown et al., 2015b). These results are consistent with an in silico study showing that the *Hic1* gene contains multiple RA receptor response elements (RAREs) (Lalevée et al., 2011). However, HIC1 is dispensable for RA-mediated expression of intestinal homing molecules and RA-dependent suppression of IL-17A production in vitro, suggesting that the RA-HIC1 axis plays a specific role in regulating intestinal T cell homeostasis that remains to be defined.

T cell-intrinsic deletion of HIC1 resulted in a significant reduction in the frequency and number of T cells in the intestinal microenvironment. Interestingly, mice raised on a VAD diet also display reduced numbers of intestinal T cells (Iwata et al., 2004). As RA has been shown to regulate expression of intestinal homing molecules such as CCR9 and α4β7 integrin, we first suspected that defective trafficking to the intestine was responsible for the paucity of T cells in the intestines of *Hic1^ΔT^* mice; however, we did not observe any defects in the ability of *Hic1*-deficient T cells to express CCR9 or α4β7. In addition, we observed a significant influx of T cells into intestinal tissues upon induction of inflammation, further suggesting that migration to the intestine is not regulated by expression of HIC1. However, we did observe that the *Hic1*-deficient T cells present in the intestine at steady state expressed significantly lower levels of the surface molecules CD69 and CD103. These molecules are expressed by tissue-resident cells and are required for retention in tissues. Indeed, similar to our results, CD103-deficient mice have a severe reduction in the number of T cells in the intestine (Schön et al., 1999). Thus, HIC1 appears to be required for the optimal expression of CD69 and CD103 and retention of T cells in the intestinal microenvironment.

In addition, CD69 and CD103 are characteristic markers of a subset of memory T cells known as tissue resident memory T (T_RM_) cells. T_RM_ cells are long-lived quiescent cells that are thought to have derived from effector T cells that have migrated into non-lymphoid tissues. Interestingly, HIC1 has been shown to control fundamental cellular processes such as cell growth and survival (Rood and Leprince, 2013; Dehennaut et al., 2012). HIC1 has been shown to interact with and regulate key transcription factors involved in cell cycle progression and cellular metabolism (Van Rechem et al., 2010; Li et al., 2012). For example, HIC1 is involved in a regulatory feed back loop with the deacteylase SIRT1 (Chen et al., 2005), which is a key regulator of fatty acid oxidation (Gerhart-Hines et al., 2007). Further, it has been demonstrated that fatty acid metabolism is key to memory T cell development and survival (van der Windt and Pearce, 2012; Pearce et al., 2009). Thus, it is possible that in addition to directly regulating T cell retention in tissues, HIC1 may also be required in T cells to promote quiescence through metabolic pathways. Future studies will aim to characterize the role for HIC1 in T cell quiescence and memory development.

Another observation was the increase in the frequency of IL-17A-producing T_H_17 cells in the intestine of *Hic1^ΔT^* mice. T_H_17 cells have been shown to play an important role in host defense to extracellular bacteria and fungi (Annunziato et al., 2015). However, they have also been described as pathogenic in multiple inflammatory diseases such as psoriasis and rheumatoid arthritis, and targeting IL-17A has proven to be an effective therapy (Papp et al., 2012; Leonardi et al., 2012; Genovese et al., 2010). However, the role of IL-17A in inflammatory bowel disease is controversial. Initial studies using the T cell transfer model of colitis provided the first evidence that IL-23-driven T_H_17 cells were pathogenic (Ahern et al., 2010). However, targeted therapies against IL-17A in Crohn’s disease have been proven ineffective (Targan et al., 2012; Hueber et al., 2012) and recent studies have shown that IL-17A is important for maintaining intestinal barrier integrity and has a protective role in regard to the development of colitis (Lee et al., 2015; Maxwell et al., 2015). Further, it is becoming more apparent that the T_H_17 cell lineage has a degree of heterogeneity with regards to pathogenicity. For example, T_H_17 cells differentiated in the presence of TGFβ are less pathogenic and produce higher levels of IL-10 while cells differentiated in the presence of IL-23 are more pathogenic and can produce IFN-γ (Lee et al., 2012; McGeachy et al., 2007). In addition, T_H_17 cells have a degree of plasticity and fate-mapping studies have shown that ‘ex-T_H_17’ cells lose their ability to produce IL-17A altogether and exclusively produce IFN-γ under certain conditions (Hirota et al., 2011). As our studies show increased production of IL-17A from T_H_17 cells and a lack of pathogenicity in multiple mouse models of intestinal inflammation, we hypothesize that in the absence of HIC1, T_H_17 cells are skewed to a more protective lineage and do not transition into pathogenic cells. The precise molecular mechanisms of how HIC1 regulates the pathogenicity of T cells remains to be fully elucidated.

In summary, we have identified the transcriptional factor HIC1 as an RA-responsive cell-intrinsic regulator of T cell function in the intestine. Through its interaction with STAT3, HIC1 limits IL-17A production by T_H_17 cells and is required for the development of intestinal inflammation. Together, these results suggest that HIC1 is an attractive therapeutic target for the treatment of inflammatory diseases of the intestine such as Crohn’s disease and potentially other diseases associated with dysregulated T_H_17 cell responses.

## METHODS

### Ethics statement

Experiments were approved by the University of British Columbia Animal Care Committee (Protocol number A15-0196) and were in accordance with the Canadian Guidelines for Animal Research.

### Mice

The generation of *Hic1*^Citrine^ mice has been described (Pospichalova et al., 2011) and the generation of *Hic1^fl/fl^* mice will be described elsewhere (manuscript in preparation). *Cd4*-Cre mice were obtained from Taconic. Animals were maintained in a specific pathogen-free environment at the Biomedical Research Centre animal facility.

### Diet Studies

Vitamin A-deficient (TD.09838) diet was purchased from Harlan Teklad Diets. At day 14.5 of gestation, pregnant females were administered the vitamin A-deficient diet and maintained on diet until weaning of litter. Upon weaning, females were returned to standard chow, whereas weanlings were maintained on special diet until use.

### Antibodies and flow cytometry

Absolute numbers of cells were determined via hemocytometer or with latex beads for LP samples. Intracellular cytokine (IC) staining was performed by stimulating cells with phorbol 12-myristate 13-acetate (PMA), ionomycin, and Brefeldin-A (Sigma) for 4 hours and fixing/permeabilizing cells using the eBioscience IC buffer kit. All antibody dilutions and cell staining were done with PBS containing 2% FCS, 1 mM EDTA, and 0.05% sodium azide. Fixable Viability Dye eFluor 506 was purchased from eBioscience to exclude dead cells from analyses. Prior to staining, samples were Fc-blocked with buffer containing anti-CD16/32 (93, eBioscience) and 1% rat serum to prevent non-specific antibody binding. Cells were stained with fluorescent conjugated anti-CD11b (M1/70; in house), anti-CD4 (GK1.5), anti-CD8 (53-6.7), anti-TCRβ (H57-597), anti-MHCII (I-A/I-E) (M5/114.15.2), anti-CD11c (N418), anti-F4/80 (BM8), anti-IL17a (17B7), anti-FOXP3 (FJK-16s), anti-B220 (RA3-6B2), anti-TCRγδ (eBioGL3), anti-CD45 (30-F11), anti-CD45RB (C363.16A) anti-α4β7 (DATK32), anti-CCR9 (eBioCW-1.2), anti-IFN-γ (XMG1.2), CD25 (PC61.5), IL-13 (eBio13A) purchased from eBioscience, anti-CD103 (M290), anti-CD69 (H1.2F3), anti-CD62L (MEL-14), anti-CD44 (IM7), anti-CD64 (X54.5/7.1.1), anti-pSTAT3 (pY705) purchased from BD Biosciences. Data were acquired on an LSR II flow cytometer (BD Biosciences) and analyzed with FlowJo software (TreeStar).

### RNA isolation and quantitative real-time PCR

Tissues were mechanically homogenized and RNA was extracted using the TRIzol method according to the manufacturer’s instructions (Ambion). cDNA was generated using High Capacity cDNA reverse transcription kits (Applied Biosystems). Quantitative PCR was performed using SYBR FAST (Kapa Biosystems) and SYBR green-optimized primer sets run on an ABI 7900 real-time PCR machine (Applied Biosystems). Cycle threshold (C_T_) values were normalized relative to beta-actin (*Actb*) gene expression. The primers used were synthesized de novo:

*Hic1:*

forward 5’-AACCTGCTAAACCTGGACCAT-3’

reverse 5’-CCACGAGGTCAGGGATCTG-3’

*Il17a*:

forward 5’-AGCAGCGATCATCCCTCAAAG-3’

reverse 5’-TCACAGAGGGATATCTATCAGGGTC-3’

*Ifng:*

forward 5’-GGATGCATTCATGAGTATTGCC-3’

reverse 5’-CCTTTTCCGCTTCCTGAGG-3’

*Il13:*

forward 5’-CCTGGCTCTTGCTTGCCTT-3’

reverse 5’-GGTCTTGTGTGATGTTGCTCA-3’

*Foxp3:*

forward 5’-CCCAGGAAAGACAGCAACCTT-3’

reverse 5’-TTCTCACAACCAGGCCACTTG-3’

*Actb:*

forward 5’-GGCTGTATTCCCCTCCATCG-3’

reverse 5’-CCAGTTGGTAACAATGCCATGT-3’.

### Cell Culture

CD4^+^ T cells were isolated from spleen and LNs by negative selection using an EasySep Mouse CD4^+^ T cell isolation kit (StemCell Technologies). 5 x 10^5^ CD4^+^ cells were cultured for up to 5 days in DMEM supplemented with 10% heat-inactivated FBS, 2 mM glutamine, 100 U/ml penicillin, 100 μg/ml streptomycin, 25 mM HEPES, and 5 x 10^−5^ 2-ME with 1 μg/ml plate bound anti-CD3 (145-2C11) and anti-CD28 (37.51). CD4^+^ T cells were polarized under T_H_17 cell-(20 ng/ml IL-6, 10 ng/ml IL-23, TNF-α, IL-1β, 1 ng/ml TGF-β1, 10 μg/ml anti-IFN-γ and anti-IL-4), T_H_1 cell-(10 ng/ml IL-2, IL-12 and 5 μg/ml anti-IL-4), T_H_2 cell-(10 ng/ml IL-2, IL-4 and 5 μg/ml anti-IFN-γ) or iT_reg_ cell-(10 ng/ml IL-2 and TGFβ1) promoting conditions. In some cases, cells were plated as above in the presence of 10 nM Retinoic Acid (Sigma-Aldrich).

## ELISA

IL-17A production was analyzed from supernatants taken on day 4 of CD4^+^ T cell *in vitro* culture using commercially available antibody pairs (eBioscience).

### Anti-CD3-induced intestinal inflammation

Mice were administered 30 μg of anti-CD3ε antibody (145-2C11) in 400μl PBS by intraperitoneal injection. Animals were monitored daily after injection and euthanized at 48 hours post injection. At endpoint, sections of the distal ileum were fixed in 10% buffered formalin, embedded in paraffin and stained with H&E. Histological inflammation were blindly scored on a scale of 0 to 4, where 0 represented a normal ileum and 4 represented severe inflammation. Specific aspects such as infiltrating immune cells, crypt length, epithelial erosion, and muscle thickness were taken into account.

### T cell Transfer Colitis

CD4^+^ cells were enriched from spleens and peripheral LNs (pLNs) of *Hic1^fl/fl^* or *Hic1^ΔT^* mice with an EasySep Mouse CD4^+^ T cell isolation kit (StemCell Technologies) and stained with anti-CD4, anti-CD25 and anti-CD45RB. Naive CD4^+^CD25^−^CD45RB^hi^ cells were purified by cell sorting. CD4^+^CD25^−^CD45RB^hi^ naive T cells (4 × 10^5^) were injected intraperitoneally into age-and sex-matched *Rag1^−/−^* mice, which were monitored for weight loss and sacrificed 6 weeks after initiation of the experiment. At endpoint, proximal colon was fixed, embedded, and stained with H&E. Histological inflammation was scored as above.

### Isolation of Intraepithelial and Lamina Propria Leukocytes

Peyer’s patches were removed from the small intestine, which was cut open longitudinally, briefly washed with ice-cold PBS and cut into 1.5 cm pieces. Tissue was incubated in 2mM EDTA PBS for 15 minutes at 37°C and extensively vortexed. Supernatants were collected and pelleted then re-suspended in 30% Percoll solution and centrifuged for 10 minutes at 1200 rpm. The pellet was collected and used as intraepithelial leukocytes. Remaining tissue was digested with Collagenase/Dispase (Roche) (0.5 mg/mL) on a shaker at 250 rpm, 37°C, for 60 minutes, extensively vortexed and filtered through a 70μm cell strainer. The flow-through cell suspension was centrifuged at 1500rpm for 5 min. The cell pellet was re-suspend in 30% Percoll solution and centrifuged for 10 minutes at 1200 rpm. The pellet was collected and used as lamina propria leukocytes.

### Cell lysis, Immunoprecipitation and Immunoblotting

Cells were lysed and immunoprecipitation carried out using antibodies against STAT3 (C-20; Santa Cruz) and FLAG (M2; Sigma). Immunoblotting was carried out using antibodies against STAT3, HIC1 (H-123; Santa Cruz) and GAPDH (GA1R; in house).

### Chromatin Immunoprecipitiation

Naive CD4^+^ T cells were activated and T_H_17 polarized for 3 days, followed by crosslinking for 8 minutes with 1% (vol/vol) formaldehyde. Cells were collected, lysed and sonicated. After being precleared with protein A agarose beads (Upstate), cell lysates were immunoprecipitated overnight at 4 °C with anti-STAT3 (C-20, Santa Cruz) or normal rabbit IgG (Cell Signaling). After washing and elution, crosslinks were reversed for 4 h at 65°C. Eluted DNA was purified and samples were analyzed by quantitative real time PCR on a 7900 Real-Time PCR system (Applied Biosystems). Primer sets used for analysis are: *Il17a* promoter forward 5’-CACCTCACACGAGGCACAAG-3’ and reverse 5’-ATGTTTGCGCGTCCTGATC-3’; *Il5* promoter forward 5’-AAGTCTAGCTACCGCCAATA-3’ and reverse 5’-AGCAAAGGTGAGTTCAATCT-3’. Each Ct value was normalized to the corresponding input value.

### Retroviral Transduction

Platinum E cells were transiently transfected using the calcium phosphate method with MigR1 expression plasmids encoding GFP alone or FLAG-HIC1. Viral supernatants were collected after 48h, supplemented with 8 μg/ml polybrene (EMD Millipore) and added to T cells that had been activated under T_H_17 cell-polarizing conditions for 48 h. T cells (2 x 10^6^) were incubated in 24 well plates with 1 ml viral supernatants. After 24 hours, viral supernatant was replaced with conditioned culture medium and cells were cultured under T_H_17 cell-polarizing conditions for an additional 3 days.

### Statistics

Data are presented as mean ± S.E.M. Statistical significance was determined by a two-tailed Student’s *t*-test using GraphPad Prism 5 software. Results were considered statistically significant with *P* < 0.05.

## AUTHOR CONTRIBUTIONS

K.B., F.A., A.C., M.B., S.S. and C.Z. designed and performed the experiments. T.M.U. and V.K. provided assistance and contributed reagents and materials. K.B. and C.Z. wrote the manuscript.

## ACKNOWLEDGMENTS

We would like to thank C. Hunter, K. Jacobson and S. Gerondakis for advice on the manuscript. We would like to thank R. Dhesi, L. Rollins (BRC core), A. Johnson (UBCFlow), M. Williams (UBC AbLab), T. Murakami (BRC Genotyping), I. Barta (BRC Histology), and all members of BRC mouse facility for excellent technical assistance. This work was supported by the Canadian Institutes of Health Research’s (CIHR) Canadian Epigenetics, Environment and Health Research Consortium (grant 128090 to C. Zaph) and operating grants (MOP-89773 and MOP-106623 to C. Zaph) and Australian National Health and Medical Research Council (NHMRC) project grants (APP1104433 and APP1104466 to C. Zaph). F. Antignano is the recipient of a CIHR/Canadian Association of Gastroenterology/Crohn’s and Colitis Foundation of Canada postdoctoral fellowship. C. Zaph is a Michael Smith Foundation for Health Research Career Investigator and a veski innovation fellow.

## FIGURES AND LEGENDS

**Figure 1-figure supplement 1.**
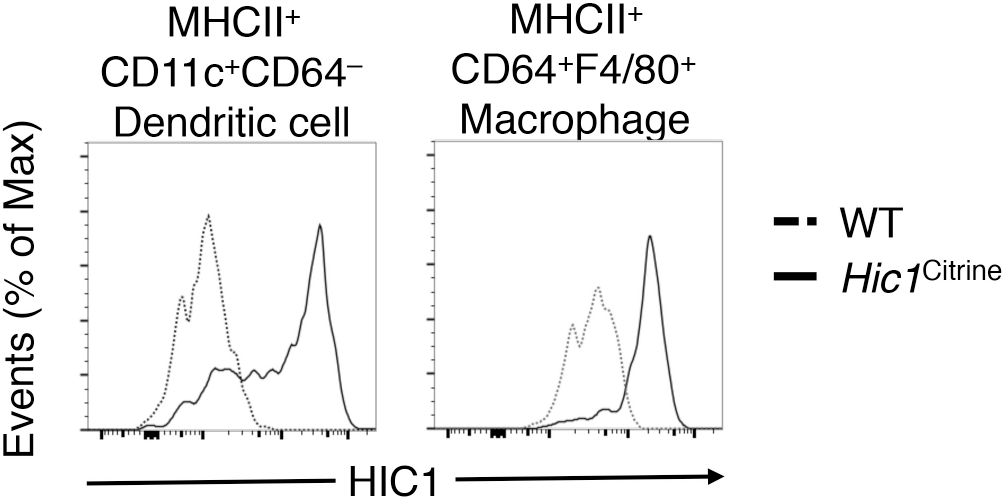
HIC1 is expressed in dendritic cells and macrophages in the intestine. MHCII^+^CD11c^+^CD64^−^ Dendritic cells and MHCII^+^CD64^+^F4/80^+^ Macrophages from intestinal lamina propria were analyzed for Hic1^Citrine^ expression by flow cytometry. Data are representative of 2 independent experiments.

**Figure 2-figure supplement 1.**
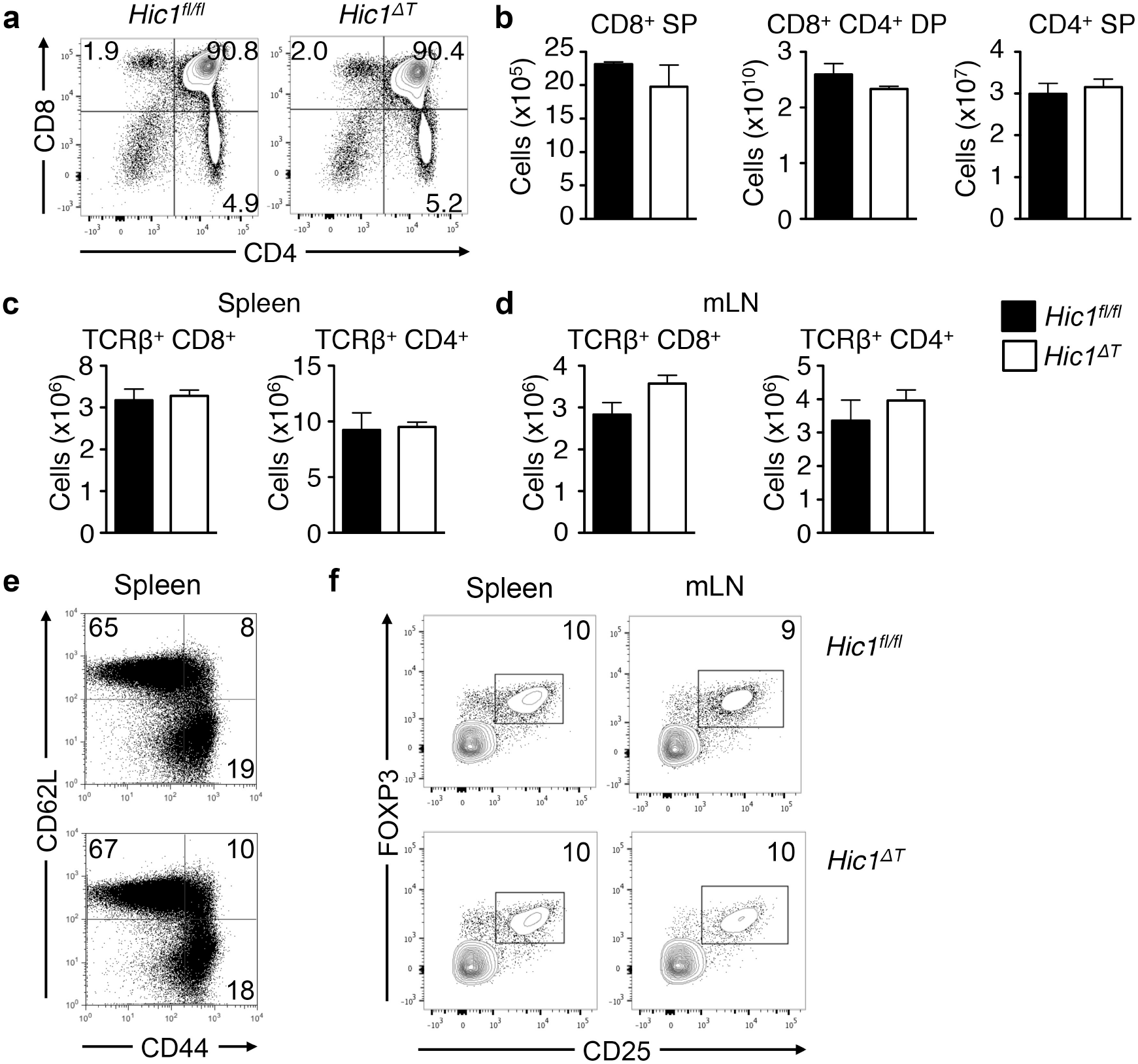
Normal central and peripheral T cell responses in the absence of HIC1. (a-d) Thymus, spleen and mesenteric lymph nodes (mLNs) from *Hic1*^fl/fl^ and *Hic1*^−/−^ mice were analyzed for CD4^+^ and CD8^+^ T cell frequencies and numbers by flow cytometry. (SP=single positive, DP=double positive). (e) CD62L^+^ and CD44^+^ frequencies of splenic CD4^+^ T cells were analyzed by flow cytometry. (f) CD25^+^FOXP3^+^ T_reg_ frequencies were analyzed from the spleen and mLN by flow cytometry. (b-d) Data are pooled from 3 independent experiments (n=4-6 per group). * *P* < 0.05. Error bars indicate SEM.

**Figure 3-figure supplement 1.**
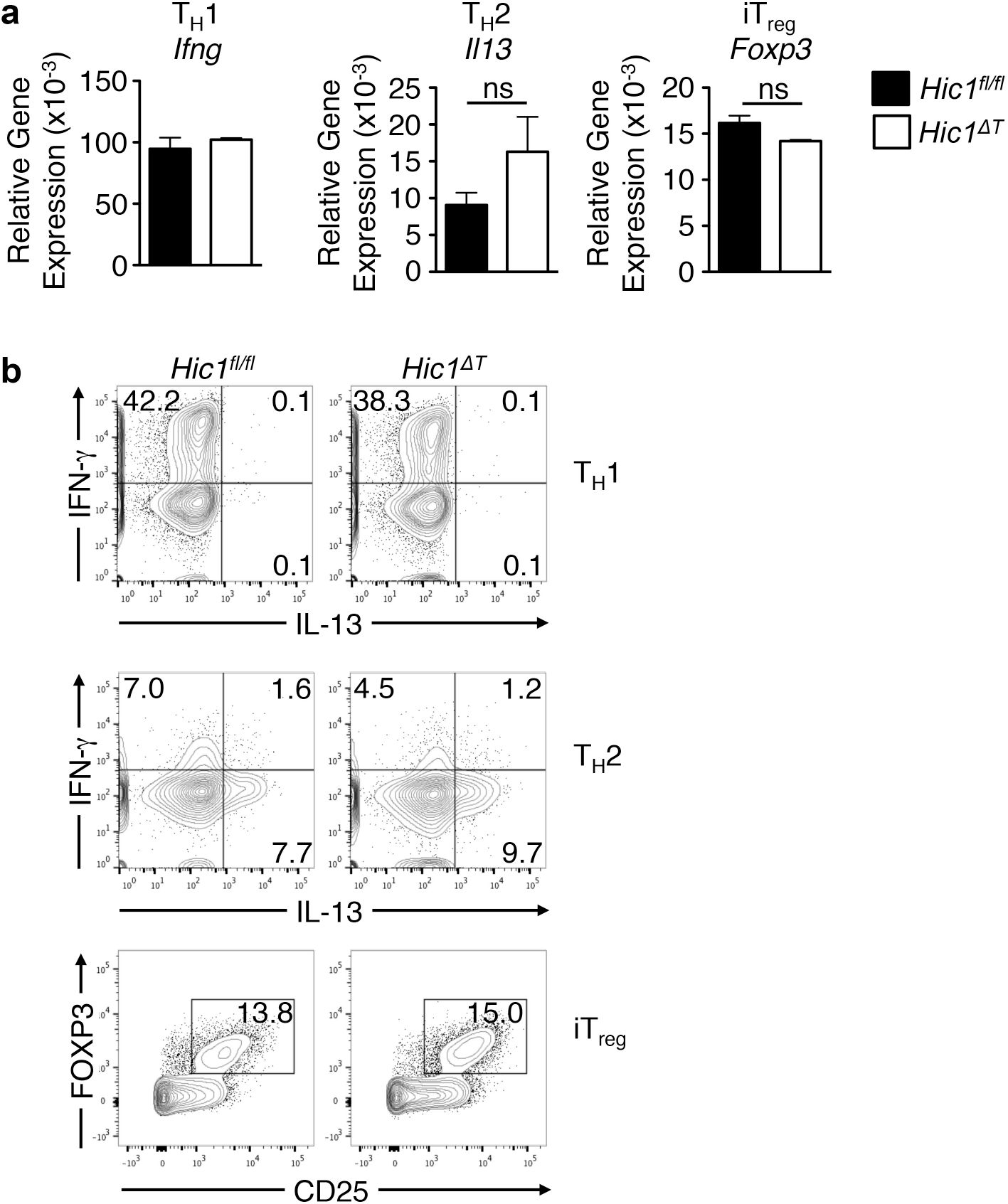
HIC1 is dispensable for T_H_1, T_H_2 and iT_reg_ cell differentiation. *Hic1^fl/fl^* or *Hic1^ΔT^* splenic CD4^+^ T cells were activated under T_H_1, T_H_2 or iT_reg_ cell-polarizing conditions and analyzed for: (a) *Ifng*, *Il13* or *Foxp3* mRNA expression by quantitative RT-PCR, and (b) intracellular IFNg, IL-13 and FOXP3 by flow cytometry. Data are pooled from 2 independent experiment (n=6 per group). * *P* < 0.05. Error bars indicate SEM.

**Figure 5-figure supplement 1.**
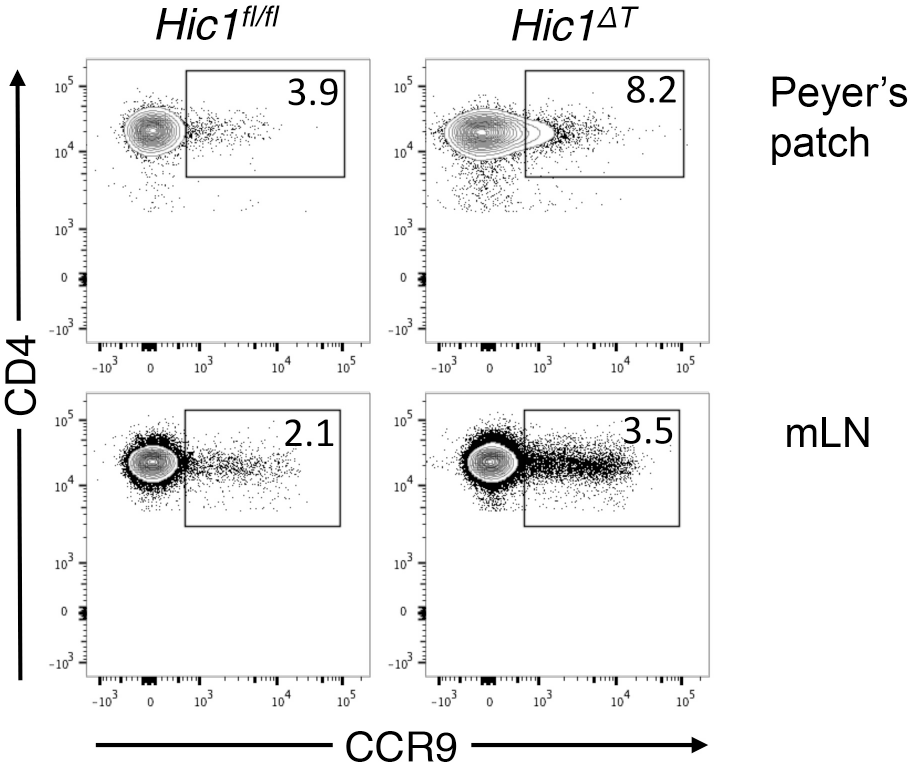
HIC1 does not regulate expression of CCR9 on CD4^+^ T cells in the intestinal lymphoid tissues. (a) CD4^+^ T cells were isolated from the mesenteric lymph nodes (mLN) and peyer’s patches (PP) and analyzed for CCR9 expression by flow cytometry. Data are representative of 2 independent experiments.

